# Development of a new bioinspired peptide with fibroblast relaxation proprieties for cosmetics applications

**DOI:** 10.1101/2023.01.27.525652

**Authors:** Márcia Renata Mortari, João Daivison Silva Ramalho, Nichollas Serafim Camargo, Guilherme Alves Ferreira, Sheila Siqueira Andrade, João Paulo Figueiró Longo

## Abstract

The cosmetic field is driven by the development of new inputs and raw materials that are used to the development innovative products for consumers. Among the different sub-areas in cosmetology, skin care is the biggest area, with almost fifty percent of the market, and probably the most innovative of them. Thus, there is a complex value chain organized to develop and produce innovative inputs to supply the demands of the market. Within this context, different basic scientific areas, such as biochemistry and biotechnology, have been a source of inspiration for new active ingredients. In the present article, we will present the pre-clinical and clinical trials involved in the development of a new innovative biomimetic peptide derived from natural wasp venoms content. This new peptide sequence, here named WASP-PEP was designed based on natural peptide templates with some strategic amino acid modifications to increase the molecule’s effectiveness. As the main results, we observed that the WASP-PEP is a biocompatible and dermatology tolerate compound, and the peptide can promote some relaxation effects in dermal fibroblasts. This relaxation effect is compatible with a kind of regulated and cumulative botox-like effect. Notablyr, this relaxation effect was also observed in the clinical trial, when volunteer subjects reported that their facial skin was flatter after the use of the WASP-PEP product for one month.

**Graphical Abstract:** 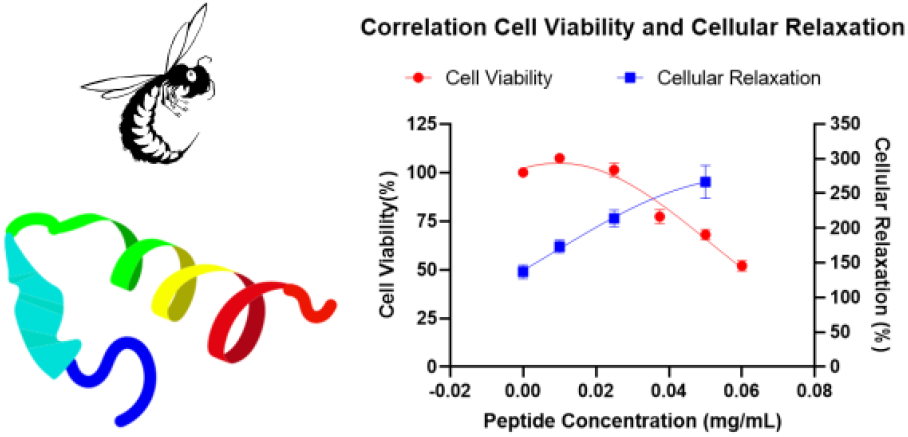

## Background

The cosmetic field is a huge market that includes all the products applied over the skin and/or hair to improve appearance or protect it against environmental stress factors. In terms of economic impact, the market has, in the last years, a continuous growth rate of 6% per year, which is an important growth profile. Regarding the cosmetic field subcategories, the largest is skincare, representing 42% of the market, with an estimative projection of worldwide revenue close to 177 billion US dollars in 2025 ^1^.

Regarding the cosmetic field organization, there are several players in the chain value, including brand developers, local sellers, global distributors, local and global industries, and cosmetic ingredient developers. This last player is represented by research (1) labs placed in universities; (2) big research and development laboratories, usually associated with big cosmetic companies; and (3) start-ups and spin-off companies that contribute to the innovation process by releasing their innovations to the market.

All these players are intrinsically connected in a complex cosmetic innovation ecosystem that creates all the novelties and new products for this growing market. In this scenario, the triggers for new developments can be activated by any of these actors, but usually, the more successful ones are derived from the combined efforts of the innovation and marketing areas. In other words, the marketing areas usually have relevant and strategic consumer information, that, in combination with the innovation sectors can develop kind of tailor-made creations that will full fill the market’s desires.

Following this playbook, several innovative cosmetic ingredients have been developed over time, including natural extracts, nanotech formulations, synthetic active ingredients, etc. An increase in the use of biomimetic/bioactive peptides in personal care products has been released over the past twenty years, as part of the efforts to slow down skin biological clock, and skin aging processes, through stimulating coordination of self-renewal at the cellular level ^2-5^. Due to the great interest in these biomimetic peptides as active cosmetic ingredients, several companies and research centers are investing efforts to develop new inputs and/or ingredients based on peptide sequence design ^6^.

Peptides are defined as amino acid sequences with a wide range of functional applications, including the pharmaceutical and cosmetic areas ^3, 4^. Nowadays it is established immeasurable potential that a carefully selected biomimetic peptide molecule might offer in cosmetic science, in particular where fine-tuneable cellular and molecular mechanisms might be used to regulate and modulated cell and consequently tissue response. In the cosmetic field, these active ingredients have attracted attention in the last few years due to the excellent clinical results observed in different types of cosmetic applications. Moreover, due to their specific target mechanisms, usually acting in some specific molecular target, these active ingredients can be used in low concentrations^7^.

The inspiration for new peptide developments is diverse. They can be developed by machine learning and artificial intelligence systems ^8, 9^ and or can be designed based on sequences that already exist in nature using molecular docking and dynamic molecular ^10, 11^. For this last one, researchers defined this approach as a nature-inspired, or biomimetic, strategy and have some advantages such as the use of natural sequences, or part of them, to design new peptide sequences. As peptide stability and shelf-life are affected by amino acid sequences, the use of naturally occurring peptide templates can accelerate the development of new active peptide sequences.

Using this biomimetic approach, our research group has invested efforts in the development of different peptides for cosmetic applications. Some examples of peptides have been proposed, including a peptide inspired in a wasp venom derivate, called Wasp-Thr6-bradykinin Oligopeptide-1-Amide (International Nomenclature Cosmetic Ingredient-37372). This peptide (WASP-PEP) design was inspired by the original peptide sequence and adapted to increase their activity, as well as skin permeation. As the main cosmetic activity, this peptide has some interesting pro-aging activities, including some relaxation effects that are similar to botox-like effects. Thus, this peptide can be used for skin care product development.

The full development of this new peptide active cosmetic ingredient involved a team with different backgrounds and expertise. The full process included sequence identification and design, pre-clinical evaluation, cosmetic formulation development, clinical trial execution, and data analysis. As the peptide was recently protected in a patent deposit, we aimed to present the results involved in the development of this new peptide sequence (WASP-PEP) for cosmetic applications. Thus, the objective of this article is to present the pre-clinical and clinical results obtained during the development of this Wasp-Thr6-bradykinin Oligopeptide-1-Amide peptide.

## Methodology

### Peptide Design and Characterization

A new peptide bioinspired from the peptides naturally present in social wasp venom was conceived and designed to modify the primary structure and induce differences in efficacy and bioavailability. After obtaining the peptide sequence, searches were made for similarities with peptides. For this, appropriate databases were used, which were Uniprot (available at http://www.uniprot.org/) and BLASTP (available at <http://blast.ncbi.nlm.nih.gov/Blast.cgi?PAGE=Proteins>). After similarities had been found, the peptide sequences were aligned using the Clustal Omega software (available at <http://www.ebi.ac.uk/Tools/msa/clustal/>). As the WASP-PEP sequence was recently patented by the Brazilian Intellectual propriety agency (INPI-*Instituto Nacional de Propriedade Intelectual*), we will not present the full molecule characterization. For more details, we should wait for the publication of the patent documents (BR 10 2022 002377 8) on the INPI website. Moreover, we also registered the WASP-PEP in two international new molecule repositories. The WASP-PEP CAS number is 2759014-86-1, and the INCI number is 37372. The Chemical Abstracts Service (CAS) is a division of the American Chemical Society, while the INCI (International Nomenclature of Cosmetic Ingredients) number is provided by the Personal Care Products Council (PCPC). These two new molecule repositories are the most important database for new compounds in the cosmetic field.

### Preclinical in vitro studies

#### Cell viability assay

For the preclinical *in vitro* studies, we used fibroblasts cells cultures to evaluate potential cytotoxicity, as well as to evaluate the relaxation effect of the WASP-PEP samples. For the experiments, the Babl/C 3T3 (clone 31) cells were used and obtained from the supplier Cell Bank of Rio de Janeiro, batch 001242. The cells were kept in sterile standard culture conditions (37°C and 5% CO_2_) with DMEM (Dulbecco’s Modified Eagle’s Medium) supplemented with 10% of fetal bovine serum (FBS; Gibco^®^ Invitrogen™, USA) and 1% of antibiotic solution (100 units/mL penicillin and 100 mg/ml streptomycin), respectively.

For the cytotoxic experiments, cells were seeded in 96-well culture plates at a cell density of 10^4^ cells/well. For peptide exposure, we prepared an initial WASP-PEP (1 mg/mL) solution, diluted in the same culture media described before. For cell exposure, serial dilutions from these initial samples were prepared and cells were exposed to the tested samples for 24 hours. For each tested concentration (0.06, 0.05, 0.0375, 0.025, and 0.01 mg/mL), 4 individual well, containing the cells, were used. We also prepared two control groups for evaluation: 1. Negative control – cells treated with culture media; 2. Positive control – cells treated with sodium lauryl sulfate to induce cell lysis.

For cell viability evaluation, we used the neutral red assay ^12^. After the WASP-PEP exposition (24 hours), the culture media was removed, cells were washed with sterile PBS, and the neutral red diluted in culture media was added (100 µL) for each well. After this step, the cells were incubated for 2 hours for dye internalization. After this period, the dye solution was removed, cells were washed with PBS, and the dye was extracted. For quantification, the neutral red that was uptake by viable cells was quantified in each well by measuring the dye’s typical absorbance (540 nm). The data were analyzed by comparing the control group (set as 100%) results.

#### Cell relaxation assay

For the relaxation of fibroblast embedded in collagen disks assay, we used the culture myoblast cells (C2C12). The cells were kept in standard culture conditions, as presented for the 3T3 cells. To evaluate the contractile activity of these cells, they were embedded in a three-dimensional collagen gel, producing a scaffold that aims to mimic the dermal tissue. The collagen gel was produced with collagen type I (3mg/mL) with a mixture of 5x DMEM, NaOH, and 1x DMEM with the cells to form the collagen disk. After the polymerization of the matrix soaked with the myoblasts, the culture medium was added according to the experimental groups. For the negative control group, a three-dimensional matrix was made without the presence of cells. All groups were performed in triplicate.

To evaluate the WASP-PEP relaxation effect, we measure the collagen disk to access the myoblast contraction activity. As the control disk does not have any contraction inhibition, these disks have a higher contraction, which can be measured by the diameter reduction. On the other side, if the WASP-PEP has relaxation effects, we can observe the collagen disks with a wider diameter, showing the contraction inhibition of the tested compound. For data analysis, we prepared a dose-response between WASP-PEP concentration and the collagen disk diameters.

#### Clinical Trial

The clinical trial included 32 volunteer women who applied to participate in the study. The clinical protocol was submitted and approved by the human ethics committee from *Plataforma Brasil* (www.plataformabrasil.saude.gov.br), an organization linked to the Brazilian Ministry of Health. The approval protocol number is 5.121.752, and all the information related to the trial can be accessed on the referred website. Previously to volunteer inclusion, all participants received and signed an informed consent document describing all the risks and benefits to participate in the clinical trial. To participate in the study, the volunteers should fulfill all the inclusion and exclusion criteria presented in Table 1.

**Table 1:** Inclusion and Exclusion Criteria for the Clinical Trial

| **Table 1: Inclusion and Exclusion Criteria for the Clinical Trial** |  |
| --- | --- |
| **Inclusion Criteria** |  |
| **1** | Age 35 to 65 years; |
| **2** | Gender: male and female |
| **3** | Presence of wrinkles on the face; |
| **4** | Be in healthy condition, corresponding to the requirements pre-defined by the doctor responsible; |
| **5** | Intact skin in the test region (face); |
| **6** | The occasional user of category products; |
| **7** | Phototype I to IV according to the Fitzpatrick classification; |
| **8** | No history of irritation and/or allergy to the material used in the study; |
| **9** | Agreement to sign the informed consent form; |
| **10** | People who want to participate in the research freely and spontaneously, without profit or financial support, with only the costs of transportation and meals being reimbursed. |
| **Exclusion Criteria** |  |
| **1** | Skin diseases: vitiligo, psoriasis, lupus, atopic dermatitis; |
| **2** | Skin marks in the experimental area that interfere with the evaluation of possible reactions, skin (pigmentation disorders, vascular malformations, scars, increased, pilosity, freckles and nevus in great quantity, sunburn); |
| **3** | Active dermatoses (local and disseminated) that may interfere with the study results; |
| **4** | Pregnant or lactating women; |
| **5** | History of allergic reactions, irritation, or intense feelings of discomfort to |
| **6** | tested product category; |
| **7** | Participants with a history of allergy to the material used in the study; |
| **8** | History of atopy; |
| **9** | People with immunodeficiencies; |
| **10** | Kidney, heart, or liver transplants; |
| **11** | Intense sun exposure or tanning session up to 15 days before the initial assessment; |
| **12** | Use of the following topical systemic medications: immunosuppressants, anti-histamines, non-steroidal anti-inflammatory drugs, and corticosteroids up to two weeks before selection; |
| **13** | Treatment with acidic vitamin A and/or its derivatives orally or topically up to 01 month before the beginning of the study; |
| **14** | Vaccination up to 03 weeks before the study; |
| **15** | Participating in another study; |
| **16** | Any condition not mentioned above that, in the opinion of the investigator, could compromise the evaluation of the study; |
| **17** | History of non-adherence or unwillingness to adhere to the study protocol; |
| **18** | Professionals directly involved in carrying out this protocol and their family members. |

For the clinical trial, patients received a serum product (30 g) containing the WASP-PEP in a final concentration of 0.01 mg/g. A full description of WASP-PEP serum components is presented in Table 2. For the product use, volunteers were instructed to collect 0.6 g of the serum with the dropper applicator (one sample) and apply it to the lateral and front of the face, focusing on regions where there are greater densities of wrinkles. Moreover, they were informed to avoid the regions close to the eyes, as well as to the mouth. The application should be performed every day, for 30 days, around 7 PM, before going to sleep without the use of other cosmetics products.

**Table 2:** Serum WASP-PEP composition*

| **Table 2: Serum WASP-PEP composition\*** |  |  |
| --- | --- | --- |
|  | **Ingredient** | **INCI or CAS number** |
| **1** | WATER | 7732-18-5 |
| **2** | GLYCERIN | 56-81-5 |
| **3** | POLOXAMER 407 | 9003-11-6 |
| **4** | S-PAPER WASP THR6- BRADYKININ OLIGOPEPTIDE-1 AMIDE | 2759014-86-1 |
| **5** | PHENOXYETHANOL | 122-99-6 |
| **6** | CAPRYLYL GLYCOL | 1117-86-8 |

\*The final concentrations will not be presented due to intellectual proprieties rights.

For the safety and effectiveness evaluations, volunteers were examined by a dermatologist on days 0 and 30. Additionally, they were also instructed to report any kind of adverse effect observed during the use of the product. Regarding the product effectiveness, volunteers were evaluated by a dermatologist for six different dermatological parameters: (1) eye region wrinkles; (2) nasolabial folds; (3) forehead wrinkles; (4) elasticity; (5) firmness; and (6) hydration. The specialist examinator evaluated these six parameters on days 0 and 30 after the use of the WASP-PEP product. The semi-quantitative evaluation was performed according to Tsukahara et al (2000) ^13^, which classified the skin improvements on a subjective scale from 0-4. The lower the result, the better the individual’s perception of each evaluated item. Moreover, after the end of the study, the volunteers were invited to answer a questionary, as presented in Table 3.

**Table 3:** Questions made (yes or no) for clinical trial volunteers after the end of the trial.

|  |  |  |
| --- | --- | --- |
|  | **Question** | **% of Positive answers** |
| **1** | Have you noticed attenuation of wrinkles and expression lines in the region where the product was applied? | 66 |
| **2** | Have you noticed attenuation of imperfections in the region where the product is applied? | 69 |
| **3** | Have you noticed a reduction in the feeling of tired skin? | 84 |
| **4** | Did you notice that your skin became healthier and more vibrant? | 78 |
| **5** | Did you notice the skin is more filled and illuminated? | 69 |
| **6** | Did you notice your skin is more hydrated? | 84 |
| **7** | Did you like the product? | 84 |
| **8** | Would you buy the product? | 78 |

## Results and Discussion

We have observed a great biotechnological advance in recent years, greatly driven by the COVID-19 pandemic situation showing that several areas of biotechnology can contribute to the development of new solutions and/or materials for most different areas ^14^. The cosmetic field is not different, and the development of new active ingredients will receive a huge effort from biotech and material science researchers, and probably several examples of biotech materials will be released in the market in the next years ^15, 16^.

Among the biotech subareas, the design of new peptides will be a relevant application, since it has the potential to produce solutions for different areas of human activity, including the cosmetic field. For the peptide evaluated in the present article, the design was conducted with a biomimetic strategy. Based on the literature, we detected that some wasp venom peptides have some interesting biological applications, including the sensibilization of some surface receptors that will promote the biological effect ^17^.

After the identification of this active molecule sequence, some amino acid modifications were proceeded to improve the peptide stabilization, as well as increase the peptide activity. Moreover, as a peptide designed for topical applications, we also included a cell penetration peptide (CCP), to improve the peptide skin permeation, thus promoting better clinical activity. It is important to note that the initial strategy is to deliver this peptide to the dermal layers to promote the relaxation of cutaneous fibroblasts ^18^. In this context, this peptide final sequence was composed of two main components: the first portion formed by bioactive peptide and the second portion formed by CPP portion. Important to highlight here that this WASP-PEP sequence was recently patented and registered as a new molecule, receiving both CAS number (2759014-86-1) and INCI identification (37372). These are the two most important repositories for new compounds and active ingredients in the cosmetic field.

Following this first peptide characterization, the next step for the project development was the pre-clinical *in vitro* investigation, to investigate both: the potential peptide cytotoxicity, and the potential peptide relaxation effect to confirm the botox-like activity. For this approach, the cell line used for these investigations was the NIH-3T3 fibroblasts, derived from embryonic mice tissues. As they are the main cells of the dermis, the use of fibroblasts in preclinical *in vitro* studies has been the most widely used strategy for investigating the biocompatibility of new cosmetic ingredients ^19^.

The first point for the *in vitro* investigation was the evaluation of the *in vitro* WASP-PEP biocompatibility. The idea was to determine the maximal non-toxic *in vitro* concentration for the next efficacy evaluations. According to Figure 1, we can observe that the WASP-PEP cytotoxicity is dose-dependent, with a lethal concentration of 50% (LC50%) of 0.06135 mg/mL. Moreover, analyzing the statistical differences among the different concentrations tested, we observed that cells treated with peptide concentrations higher than 0.0375 mg/mL presented cell viability reduction, with a statistical difference compared to the control (0 mg/mL) cells. And, on the opposite, cells treated with the WASP-PEP in 0.01 and 0.025 mg/mL presented cell viability with no statistical difference (p<0.05) in comparison to the control cells.

**Figure 1:**
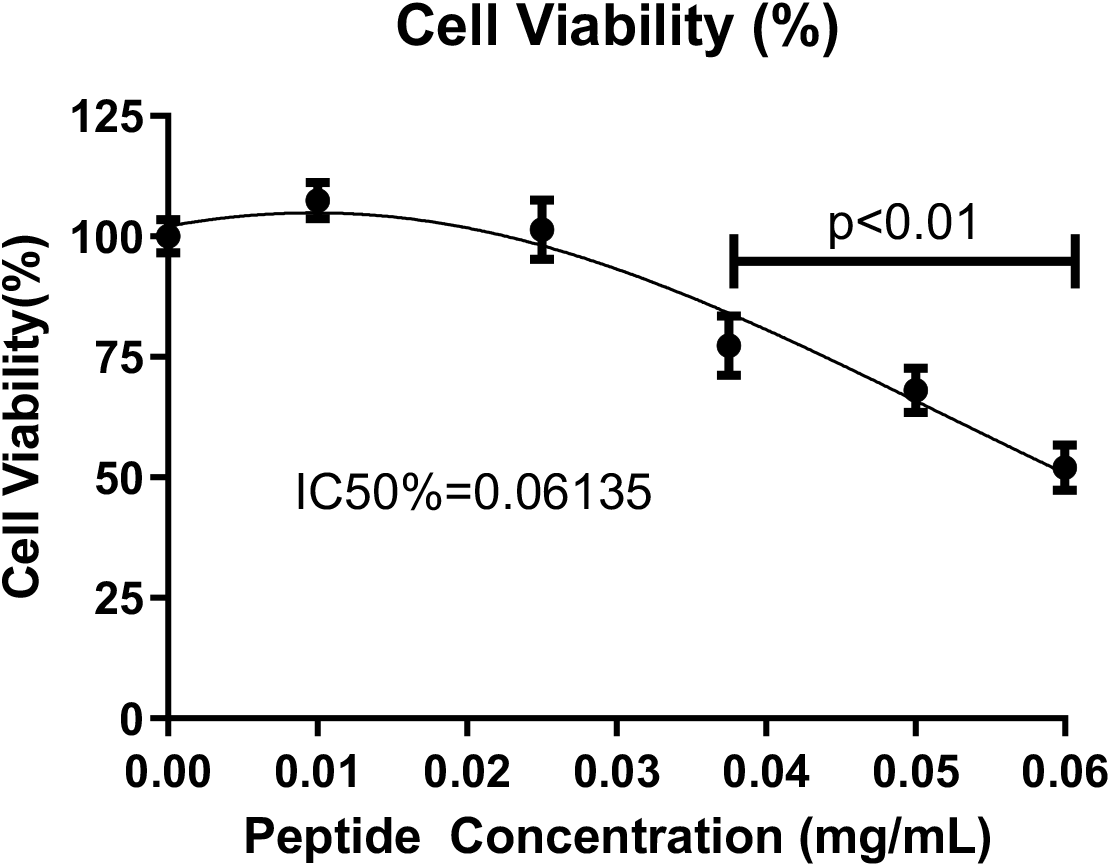
Cell viability measured by the neutral red cell cytotoxicity assay. The inhibition concentration for 50% (IC50%) was also calculated. P<0.01 represent the statistical significance of this analysis.

This kind of result is very common in the literature and reflects the dose-dependent effect of a tested compound. Higher concentrations of peptides especially with CPP sequences can increase the biocompatibility profile in higher concentrations. This result is quite expected, since the CPP sequences interact with cell membranes and promote the disruption of these structures, promoting cell death ^20^. Thus, is very important to evaluate this tolerance concentration to prevent any kind of undesirable side effects during peptide exposition. Thus, is fundamental to proceed with these pre-clinical biocompatible assays to ensure the safe concentration that should be used in the clinical evaluations.

The next step during the *in vitro* investigation was to access the botox-like proprieties. This potential activity was ideated during the WASP-PEP design and is based on the potential peptide mechanism, which includes the interaction of the peptide structure with specific fibroblast receptors. In a study using a portion of the new peptide, the authors proved that after intracerebroventricularly injection, Thr6-BK acts through B2 bradykinin receptors in the mammalian CNS, evoking analgesic behavior. This activity is remarkably different from that of bradykinin, despite the structural similarities between both peptides ^17^. Moreover, the ancestral peptide that formed a portion of the new peptide causes irreversible paralysis in insects via the presynaptic blockade of cholinergic transmission induced by the non-competitive inhibition of choline uptake, similar to the effect of hemicholinium ^21, 22^. As the WASP-PEP is a new molecule, and this relaxation/botox-like effect was not been evaluated before, this part of the investigation was fundamental to prove the effectiveness of the designed sequence.

The method used was the evaluation of the *in vitro* fibroblast contraction. This method has been widely used for different types of skin-mimetic studies and is prepared by the production of a collagen disk embedded with fibroblasts in cell culture conditions ^23, 24^. This fibroblast–collagen-matrix model is a unique strategy to evaluate cell contraction in an environment like the natural dermal extra-cellular matrix. Since fibroblasts are always anchored to the extracellular matrix, this protocol represents a close condition representing the natural conditions of dermal tissues. Moreover, as collagen fibers are one of the main components of the extracellular matrix, this methodology is an interesting approach to studying fibroblast behavior ^24^.

For the fibroblast contraction study, we used three different WASP-PEP concentrations (0.01, 0.025, and 0,05 mg/mL). The results are placed in Figure 2, and represent the relaxation effect of the peptide. As fibroblasts perform cell contraction constantly, the evaluation of their relaxation indicates that the active tested can temporarily inhibit this contractile activity. For this experiment setup, the collagen disc diameters are measured and compared among the control and peptide-exposed groups (Figure 2A). The botox-like effect is observed when the fibroblast relaxation is measured. In other words, the greater the diameter of the disc, the greater the level of cellular relaxation, that is, the lower the contractile activity of the cells embedded in the collagen discs.

**Figure 2:**
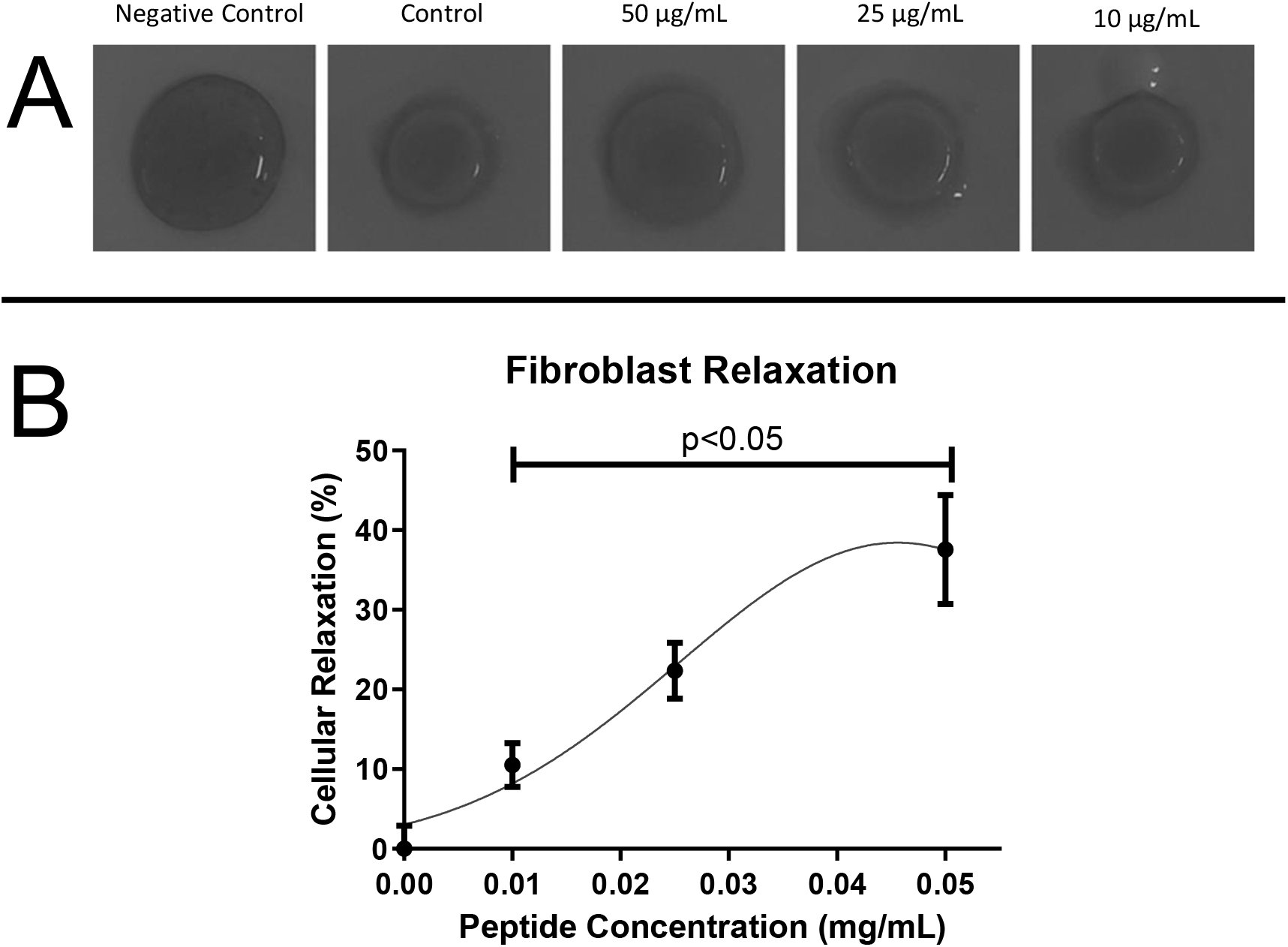
Percentage of cell relaxation, measured in collagen-embedded discs with fibroblasts. The Measurements were done using the disk’s diameters. Section A represents the collagen embedded with fibroblast diameters submitted to different experimental treatments. In section B we observe the quantification of the results obtained in the measurements of the diameter. P<0.05 represent the statistical significance of this analysis.

Regarding the relaxation results, based on Figure 2B, we can observe that the fibroblast contraction has a concentration-response effect. It shows that WASP-PEP exposure can promote fibroblast relaxation, and can potentially promote this effect in clinical conditions. In terms of fibroblast physiology, it is important to highlight here that contractile activity is typical and natural for this type of cell. As Fibroblasts are naturally involved in several physiological healing processes, they need this contractile propriety to connect the tissue elements to promote tissue regeneration. For instance, during skin wound healing, these cells can migrate and connect the scar tissue margins by using their contractile ability ^25^.

Regarding the statistical differences, we observed significance (p<0.05) for the 0.025, and 0.05 mg/mL peptide exposure. As the 0.05 mg/mL concentration presented a slight cytotoxicity effect, as shown in Figure 2B, part of this relaxation effect could be related to the cytotoxic effect. Thus, as the 0.025 mg/mL concentration promoted a significant contraction relaxation, and the cytotoxic effect was not significant, we selected this concentration to go forward in the clinical trial.

The clinical trial performed with the WASP-PEP was submitted and approved by the Human Ethics Committee (www.plataformabrasil.saude.gov.br), an organization linked to the Brazilian Ministry of Health. The approval protocol number is 5.121.752, and all the information related to the trial can be accessed on the referred website. The trial involved 32 women volunteers, with ages ranging from 37 to 49 years old. For the study, we prepared a cosmetic formulation composed of the WASP-PEP molecule entrapped in a polymer nanocarrier to increase product stability. The prototype was used as a topical serum with a defined formulation, previously described in the methodology sections. The volunteers used the serum (∼0.6 g) on the lateral face region for 30 days. The main objective of the study was to evaluate the safety and effectiveness of the serum containing the peptide after this period of use.

For the safety analysis, clinical researchers, composed of dermatologists were in charge to ask for any potential adverse effects during this one month of use. Regarding this point, no side effect or undesirable event was notified by any volunteer. This evidence confirms the clinical compatibility of the tested product. Moreover, this trial guarantees the dermatologically tested label for this new WASP-PEP serum.

The second part of the trial involved the effectiveness of the tested product. For this point, the volunteers were examined by a dermatologist on day 0 and day 30 for subjective visual perception of six different parameters: (1) eye region wrinkles; (2) nasolabial folds; (3) forehead wrinkles; (4) elasticity; (5) firmness; and (6) hydration. The results are presented in Figure 3, where the visual appearance of facial wrinkles is plotted. For all the parameters analyzed a statistically significant reduction (p<0.05) in this subjective perception was detected for all the points, except for the eye region wrinkles.

Moreover, the volunteers were also invited to answer a standard survey with several points regarding subjective evaluation and cosmetical and dermatological perception. The survey was conducted on days 0 and 30 of the clinical trial. The results obtained are placed in Table 3 and indicate an overall positive perception after these 30 days of exposure. A positive perception was detected in all questions. In terms of answer quantification, the percentage of positive responses is placed in Table 3, right column.

The clinical results described here demonstrate that the WASP-PEP evaluated has an important beneficial effect on the skin, in terms of cosmetical analysis. These results may be related to the fibroblast relaxation results described in the *in vitro* investigations. As commented before, these relaxation proprieties are also known as botox-like activities and are interesting for cosmeceutical products. Moreover, as discussed before, fibroblast cells have a contractile phenotype that is observed in normal hemostatic conditions but increases over time ^26^. Older fibroblasts have stronger contraction capacity in comparison to younger fibroblasts. This is an interesting observation, since, if we compare it with muscle cell physiology, there is a significant contraction reduction over time ^27, 28^.

For the cosmetic field, this is quite interesting, since pro-aging products are among the most desired products by consumers and by industries in the sector. Furthermore, in comparison to regular botox applications, cosmetic products with botox-like effects are topically applied, and no needles are necessary to obtain the cosmetic effect. And, comparing again the biological effects of these two products, the injected botox targets muscle cells, or even neural-muscular interface, which can promote undesirable effects for longer periods, since these tissues regenerate slowly in comparison to dermal structures ^29^.

Within this context, some initiatives like the WASP-PEP have already been proposed in the literature. For instance, the identification of some specific peptides obtained from viper venoms triggered the development of cosmetic creams with botox-like effects. The identification of neurotoxic peptides that can reduce cell contraction is the basis of all these kinds of development. In this specific case, these peptides identified from viper venoms were used as templates for the development of new innovative active cosmetic ingredients with a focus on wrinkles relaxation, which were named botox-like effects ^30^. Thus, the development of the WASP-PEP, as presented in this report, is a strategy that follows this innovation trend, that use nature as an inspiration for new compound development. For us, this is an interesting approach since it uses all the evolutionary processes developed by nature, to provide specific benefits for human beings ^10^.

On the other side, topically applied botox-like compounds, such as the WASP-PEP used in the clinical trial, probably have their activity on fibroblast cells, inducing the described relaxation effect. In terms of clinical observation, this relaxation effect is perceived as a wrinkle attenuation, since the skin will present a flatter appearance when the dermal tissues are less contracted. The clinical observation of this effect can be noted in Figure 3, where a wrinkle attenuation can be observed after the use of the WASP-PEP serum for 30 days.

## Conclusion

We describe the pre-clinical and clinical development of an innovative new peptide as an active cosmetic ingredient. The WASP-PEP described here was designed based on natural peptide templates derived from social wasp venoms, and this new peptide has some strategic amino acid modifications that aimed to increase the compound’s effectiveness. In conclusion, we observed that WASP-PEP is a dermatology tolerate active ingredient and it has some interesting cosmetic effects such as wrinkles flattening and dermal relaxation. Thus, we believe that this new biotechnology compound will be useful for the development of innovative cosmetic products.

## References

(1) Aguilar-Toalá, J. E.; Hernández-Mendoza, A.; González-Córdova, A. F.; Vallejo-Cordoba, B.; Liceaga, A. M. Potential role of natural bioactive peptides for development of cosmeceutical skin products. Peptides 2019, 122, 170170. DOI: 10.1016/j.peptides.2019.170170.

(2) Skibska, A.; Perlikowska, R. Signal Peptides-Promising Ingredients in Cosmetics. Current Protein and Peptide Science 2021, 22 (10), 716–728.

(3) Schagen, S. K. Topical peptide treatments with effective anti-aging results. Cosmetics 2017, 4 (2), 16.

(4) Mentel, M.; Schild, J.; Maczkiewitz, U.; Koehler, T.; Farwick, M. Innovative peptide technologies for even, young and healthy looking skin. SOFW Journal-Seifen Ole Fette Wachse 2012, 138 (3), 22.

(5) Blanes-Mira, C.; Clemente, J.; Jodas, G.; Gil, A.; Fernández-Ballester, G.; Ponsati, B.; Gutierrez, L.; Pérez-Payá, E.; Ferrer-Montiel, A. A synthetic hexapeptide (Argireline) with antiwrinkle activity. Int J Cosmet Sci 2002, 24 (5), 303–310. DOI: 10.1046/j.1467-2494.2002.00153.x.

(6) Usui, K.; Tomizaki, K.-y. Advances of Peptide Engineering. MDPI: 2021; Vol. 9, p 1096.

(7) Bassino, E.; Zanardi, A.; Gasparri, F.; Munaron, L. M. Effects Of The Biomimetic Peptide Sh-Polipeptide 9 (CG-VEGF) On Cocultures Of Human Hair Follicle Dermal Papilla Cells And Microvascular Endothelial Cells. 2016.

(8) Wu, Q.; Ke, H.; Li, D.; Wang, Q.; Fang, J.; Zhou, J. Recent progress in machine learning-based prediction of peptide activity for drug discovery. Current topics in medicinal chemistry 2019, 19 (1), 4–16.

(9) Gupta, R.; Srivastava, D.; Sahu, M.; Tiwari, S.; Ambasta, R. K.; Kumar, P. Artificial intelligence to deep learning: machine intelligence approach for drug discovery. Molecular Diversity 2021, 25 (3), 1315–1360.

(10) Levin, A.; Hakala, T. A.; Schnaider, L.; Bernardes, G. J.; Gazit, E.; Knowles, T. P. Biomimetic peptide self-assembly for functional materials. Nature Reviews Chemistry 2020, 4 (11), 615–634.

(11) Lima, T. N.; Pedriali Moraes, C. A. Bioactive Peptides: Applications and Relevance for Cosmeceuticals. Cosmetics 2018, 5 (1), 21.

(12) Repetto, G.; Del Peso, A.; Zurita, J. L. Neutral red uptake assay for the estimation of cell viability/cytotoxicity. Nature protocols 2008, 3 (7), 1125–1131.

(13) Tsukahara, K. A photographic scale for the assessment of human facial wrinkles. J Cosmet Sci 2000, 51, 127–139.

(14) Figueiró Longo, J. P.; Muehlmann, L. A. How has nanomedical innovation contributed to the COVID-19 vaccine development? Nanomedicine (Lond) 2021, 16 (14), 1179–1181. DOI: 10.2217/nnm-2021-0035.

(15) Ferreira, M. S.; Magalhães, M. C.; Sousa-Lobo, J. M.; Almeida, I. F. Trending Anti-Aging Peptides. Cosmetics 2020, 7 (4), 91.

(16) Longo, J. P. F.; Mussi, S.; Azevedo, R. B.; Muehlmann, L. A. Issues affecting nanomedicines on the way from the bench to the market. J Mater Chem B 2020, 8 (47), 10681–10685. DOI: 10.1039/d0tb02180f.

(17) Mortari, M.; Cunha, A.; Carolino, R.; Coutinho-Netto, J.; Tomaz, J. C.; Lopes, N. P.; Coimbra, N. C.; Dos Santos, W. Inhibition of acute nociceptive responses in rats after icv injection of Thr6- bradykinin, isolated from the venom of the social wasp, Polybia occidentalis. British journal of pharmacology 2007, 151 (6), 860–869.

(18) Cerrato, C. P.; Langel, Ü. An update on cell-penetrating peptides with intracellular organelle targeting. Expert Opinion on Drug Delivery 2022, 19 (2), 133–146.

(19) Jirová, D.; Kejlova, K.; Brabec, M.; Bendova, H.; Kolářová, H. The benefits of the 3T3 NRU test in the safety assessment of cosmetics: long-term experience from pre-marketing testing in the Czech Republic. Toxicology in vitro 2003, 17 (5-6), 791–796.

(20) Saar, K.; Lindgren, M.; Hansen, M.; Eiríksdóttir, E.; Jiang, Y.; Rosenthal-Aizman, K.; Sassian, M.; Langel, Ü. Cell-penetrating peptides: a comparative membrane toxicity study. Analytical biochemistry 2005, 345 (1), 55–65.

(21) Piek, T. Neurotoxic kinins from wasp and ant venoms. Toxicon 1991, 29 (2), 139–149.

(22) Piek, T.; Hue, B.; Mantel, P.; Nakajima, T.; Pelhate, M.; Yasuhara, T. Threonine6-bradykinin in the venom of the wasp Colpa interrupta (F.) presynaptically blocks nicotinic synaptic transmission in the insect CNS. Comparative biochemistry and physiology part C: comparative pharmacology 1990, 96 (1), 157–162.

(23) Grinnell, F. Fibroblast–collagen-matrix contraction: growth-factor signalling and mechanical loading. Trends in cell biology 2000, 10 (9), 362–365.

(24) Grinnell, F. Fibroblast biology in three-dimensional collagen matrices. Trends in cell biology 2003, 13 (5), 264–269.

(25) Darby, I. A.; Hewitson, T. D. Fibroblast differentiation in wound healing and fibrosis. International review of cytology 2007, 257, 143–179.

(26) Honeybrook, A.; Lee, W.; Woodward, J.; Woodard, C. Botulinum Toxin-A and Scar Reduction: A Review. The American Journal of Cosmetic Surgery 2018, 35 (4), 165–176.

(27) Yu, Z.; Smith, M. J.; Siow, R. C.; Liu, K.-K. Ageing modulates human dermal fibroblast contractility: Quantification using nano-biomechanical testing. Biochimica et Biophysica Acta (BBA)-Molecular Cell Research 2021, 1868 (5), 118972.

(28) Wanitphakdeedecha, R.; Kaewkes, A.; Ungaksornpairote, C.; Limsaengurai, S.; Panich, U.; Manuskiatti, W. The effect of botulinum toxin type A in different dilution on the contraction of fibroblast—in vitro study. Journal of Cosmetic Dermatology 2019, 18 (5), 1215–1223.

(29) Yue, S.; Ju, M.; Su, Z. A Systematic Review And Meta-Analysis: Botulinum Toxin A Effect on Postoperative Facial Scar Prevention. Aesthetic Plast Surg 2022, 46 (1), 395–405. DOI: 10.1007/s00266-021-02596-7.

(30) Debono, J.; Xie, B.; Violette, A.; Fourmy, R.; Jaeger, M.; Fry, B. G. Viper Venom Botox: The Molecular Origin and Evolution of the Waglerin Peptides Used in Anti-Wrinkle Skin Cream. J Mol Evol 2017, 84 (1), 8–11. DOI: 10.1007/s00239-016-9764-6.

